# A segregating human allele of *SPO11* modeled in mice disrupts timing and amounts of meiotic recombination, causing oligospermia and a decreased ovarian reserve

**DOI:** 10.1101/592576

**Authors:** Tina N. Tran, John C. Schimenti

**Affiliations:** Cornell University, Department of Biomedical Sciences and the Department of Molecular Biology and Genetics

## Abstract

A major challenge in medical genetics is to characterize variants of unknown significance (VUS), so as to better understand underlying causes of disease and design customized treatments. Infertility has presented an especially difficult challenge with respect to not only determining if a given patient has a genetic basis, but also to identify the causative genetic factor(s). Though genome sequencing can identify candidate variants, in silico predictions of causation are not always sufficiently reliable so as to be actionable. Thus, experimental validation is crucial. Here, we describe the phenotype of mice containing a nonsynonymous (proline-to-threonine at position 306) change in *Spo11*, corresponding to human SNP rs185545661. SPO11 is a topoisomerase-like protein that is essential for meiosis because it induces DNA double stranded breaks (DSBs) that stimulate pairing and recombination of homologous chromosomes.

Although both male and female *Spo11^P306T/P306T^* mice were fertile, they had reduced sperm and oocytes, respectively. Spermatocyte chromosomes exhibited synapsis defects (especially between the X and Y chromosomes), elevated apoptotic cells, persistent markers of DSBs, and most importantly, fewer Type 1 crossovers that causes some chromosomes to have none. *Spo11^P306T/−^* mice were sterile and made fewer meiotic DSBs than *Spo11^+/−^* animals, suggesting that the *Spo11^P306T^* allele is a hypomorph and likely is delayed in making sufficient DSBs in a timely fashion. If the consequences are recapitulated in humans, it would predict phenotypes of premature ovarian failure, reduced sperm counts, and possible increased number of aneuploid gametes. These results emphasize the importance of deep phenotyping in order to accurately assess the impact of VUSs in reproduction genes.

## INTRODUCTION

Properly regulated DSB formation is essential for meiotic recombination-driven pairing of homologous chromosomes during meiosis in most eukaryotic species.

SPO11, the ortholog of subunit A of Archaean topoisomerase VI, and its binding partner TOPOVIBL, work together in an evolutionarily conserved topoisomerase-like enzyme complex to generate DSBs (Robert et al. 2016; Romanienko and Camerini-Otero 2000; Baudat et al. 2000b; Vrielynck et al. 2016). In addition to this complex, SPO11 requires a partially conserved group of auxiliary proteins for chromatin recruitment and DSB-forming activity and processing. In mice, these include IHO1, MEI4, REC114, and ANKRD31 (Stanzione et al. 2016; Kumar et al. 2010; Kumar et al. 2015; Kumar et al. 2018; Boekhout et al. 2018).

*Spo11^−/−^* males are sterile and produce no sperm. They exhibit complete spermatocyte arrest in late zygonema due to severely defective homologous chromosome synapsis (Romanienko and Camerini-Otero 2000; Baudat et al. 2000b; Metzler-Guillemain and de Massy 2000). In females, loss of *Spo11* permits a subset of oocytes to progress to the dictyate stage despite being asynaptic, and actually undergo folliculogenesis. However, most are lost during prophase I from accumulation of spontaneous DSBs that trigger the DNA damage checkpoint (Rinaldi et al. 2017), and the remainder cannot fully mature or complete the first meiotic division (Romanienko and Camerini-Otero 2000; Baudat et al. 2000b; Metzler-Guillemain and de Massy 2000).

Since *Spo11* deficiency causes infertility in mice and is conserved from yeast to humans, mutations in this gene are candidates for being involved in human infertility.

There have been studies that have implicated mutations or variants in SPO11 in men with non-obstructive azoospermia (NOA) (Fakhro et al. 2018; Ren et al. 2017; Ghalkhani et al. 2014), but in no cases has causation been proven experimentally or with unassailable pedigree data. Lack of experimental validation has similarly plagued many studies of various meiosis genes and their potential roles in human infertility. This is understandable, given the absence of effective culture system to recapitulate meiosis in vitro and test the effects of particular VUSs.

We have been addressing this issue of functional validation of putative human infertility alleles using a combination of *in silico* predictions and *in vivo* modeling in mice (Singh and Schimenti 2015; Tran and Schimenti 2018). The goal is to create a permanent, reliable database of functional consequences of non-synonymous single nucleotide polymorphisms (nsSNPs) in known fertility genes (including genes that cause infertility when knocked out in mice) that will assist future genetic diagnosis of patients that are found to have one of these segregating polymorphisms in the population. Here, we report the generation and analysis of mice bearing a nsSNP (rs185545661) in the *Spo11* gene that is essential for meiosis in mice. We find that this allele indeed impacts oogenesis and spermatogenesis, and may also predispose to aneuploid gametes. Our analyses also contribute to our understanding of how formation of meiotic DSBs is regulated throughout prophase I, and emphasize the importance of timely execution of DSB formation and repair for generating crossovers on all chromosomes.

## RESULTS

### Selection of SNP rs185545661

Several criteria were considered when selecting nsSNPs as candidate infertility alleles, and ultimately modeling them in mice. First, the nsSNP must alter an amino acid conserved between humans and mice. Second, the SNP should be present at a frequency in at least one population that is not so high as to be implausible to persist in the population, but not so low as to be of trivial significance. We aimed for 0.001-2%. Third, the nsSNP must be computationally predicted to be deleterious to protein function. rs185545661 encodes a proline-to-threonine change at amino acid position 306 of human and mouse SPO11 (*SPO11^P306T^*). The P306 amino acid resides in the protein’s catalytic TOPRIM (topoisomerase-primase) domain and is highly conserved from yeast to mammals and across other related proteins with TOPRIM domains (Robert et al. 2016; Altschul et al. 1990). The *SPO11^P306T^* allele occurs at 0.0037% in the Latino population (gnomAD). The widely used SIFT and PolyPhen-2 algorithms (Kumar et al. 2009; Adzhubei et al. 2013) predict this variant to be highly deleterious to protein function, each having the “worst” possible scores of 0 and 1, respectively. Other algorithms made similar predictions, including PROVEAN (“deleterious” score = −7.61), Mutation Assessor (“high functional impact” score = 3.58), and the ensemble algorithms CADD (score = 24, in top 0.6% if all deleterious variants) and REVEL (0.517; ~10% likely of being neutral) (Calabrese et al. 2009; Ioannidis et al. 2016; Kircher et al. 2014; Choi and Chan 2015; Reva et al. 2007).

### Spo11^P306T/P306T^ mice have oligospermia and a reduced ovarian reserve

To model the Pro>Thr change encoded by rs185545661, CRISPR/Cas9-mediated genome editing was employed (Fig.1a,b). A single-strand oligodeoxynucleotide (ssODN) was used as a homologous recombination template to introduce the change. Founder mice with the correct mutation (Fig. 1C) were backcrossed into strain FVB/NJ for two generations, then intercrossed to produce homozygotes and controls for analysis.

**Figure 1.**
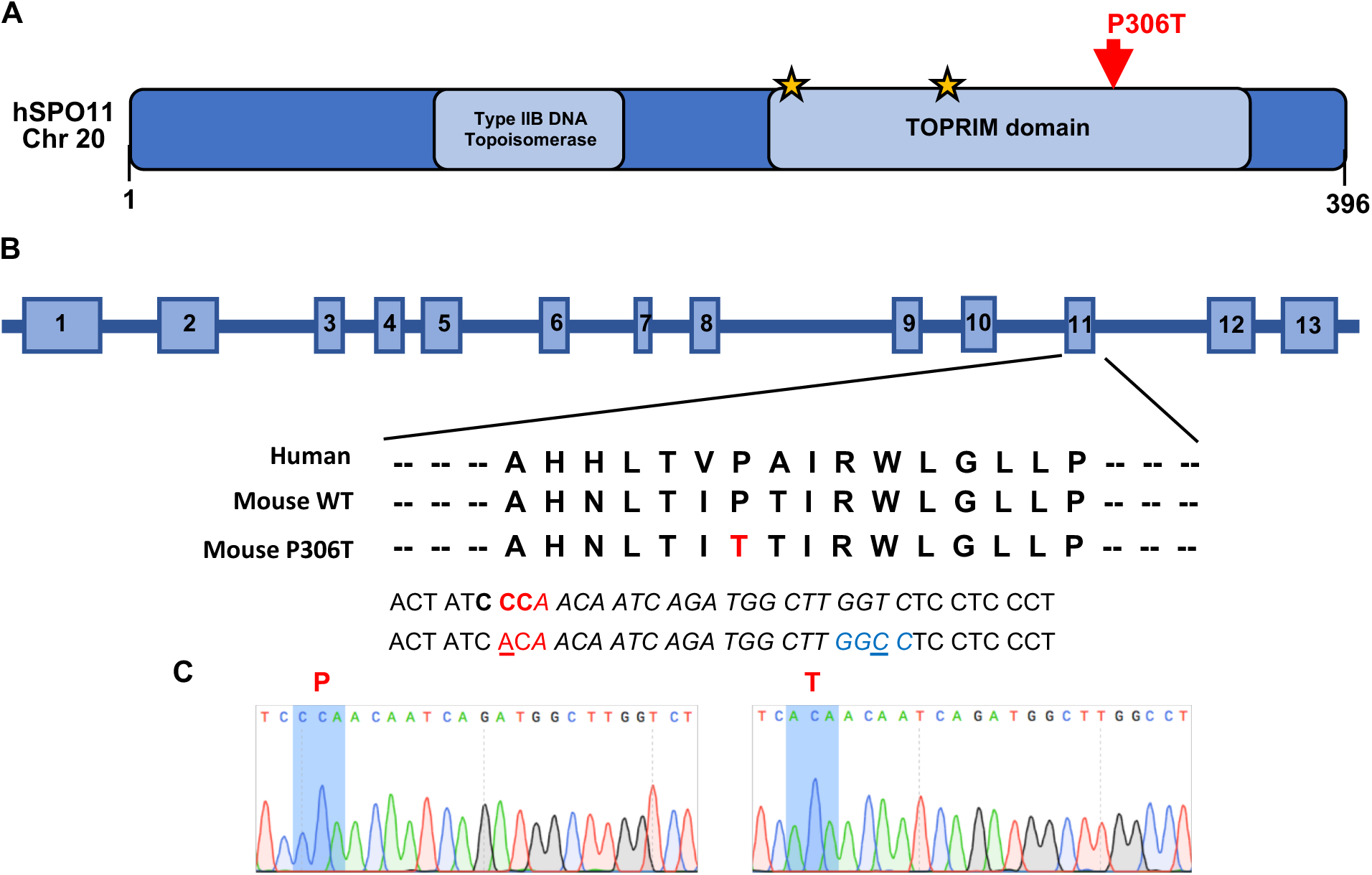
SPO11 structure and CRISPR-Cas9 editing strategy. **(A)** Schematic of human SPO11 protein with known domains and location of the P306T targeted alteration. Stars indicate known metal binding sites. **(B)** Organization of human *SPO11*. The *SPO11 P306T* variant encoded by SNP rs185545661 is located in exon 11. Red text indicates the codon encoding P306 and T306, blue text indicates a designed silent HaeIII restriction enzyme site added to the mouse allele via editing, bold text (CCC) indicates the PAM site, italicized sequence corresponds to guide sequence, and underlined text indicates nucleotide changes introduced by planned editing. **(C)** Sanger sequencing chromatograms of WT and *Spo11^P306T/P306T^* mice, with the relevant codon bases shaded blue.

To assess fertility and fecundity, adult *Spo11^P306T/P306T^* animals were bred for up to eight months with wild type (WT, or +/+) partners. Mutants of both sexes had average litter sizes that were not significantly different from WT (Fig. 2a). However, they exhibited gonadal abnormalities. Compared to WT, 8 week old *Spo11^P306T/P306T^* males had ~50% smaller testes and produced ~5 fold fewer sperm (Fig. 2b,c). Testis histology revealed that mutant seminiferous tubules supported superficially normal spermatogenesis, except that tubule diameters appeared smaller and there were substantial numbers of pyknotic-appearing spermatocytes with disorganized metaphase plates and scattered or lagging chromosomes or chromatin fragments (Fig. 2d). Consistent with the histology, mutant testes had >6 times more seminiferous tubules with >5 apoptotic cells than WT (Fig. 2e).

**Figure 2.**
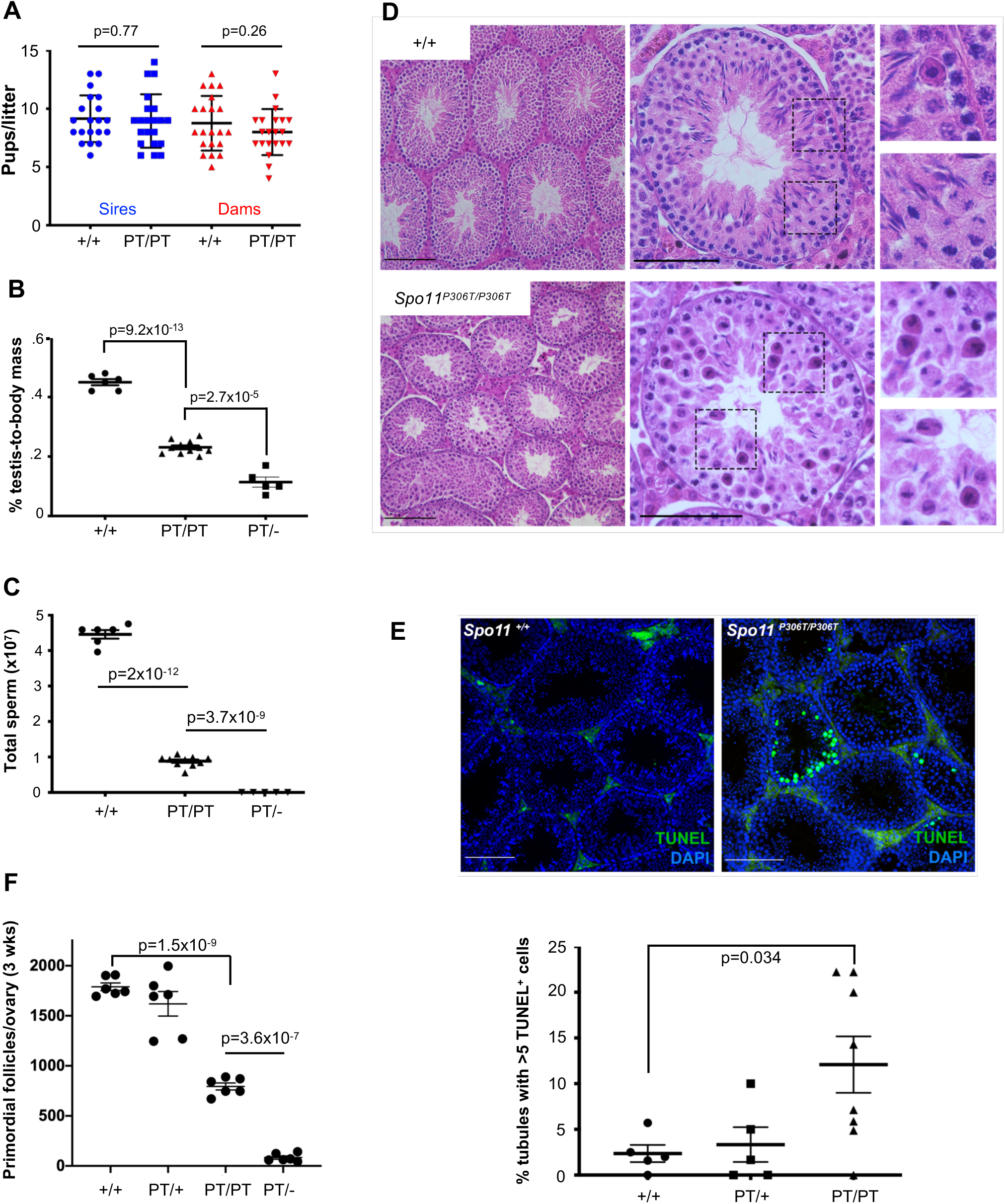
*Spo11^P306T/P306T^* mice are fertile, but have fewer sperm and a reduced ovarian reserve. **(A)** Litter sizes from matings of 2-9 month old *Spo11^+/+^* (+/+) and *Spo11^P306T/P306T^* (PT/PT) males (n=3 and 4, respectively) and females (n=3 and 4, respectively) red) to WT partners. Average litter sizes (± SEM) produced by WT and mutant males (blue) were 9.2 ± 0.45 and 9.0 ± 0.5, respectively, and from WT and mutant females (red) were 8.8 ± 0.51 and 8 ± 0.44, respectively. **(B)** Percentage of testis mass relative to body mass of 2 month old males (each data point is one animal, using average weight of both testicles). Averages were 0.45 ± 0.01, 0.42 ± 0.021, 0.23 ± 0.006 and 0.11 ± 0.0045 for each genotype from left to right. **(C)** Sperm counts from cauda epididymides of 2-month old males. Averages from left to right were 43×10^6^ ± 3.5×10^6^, 41×10^6^ ± 2.8×10^6^, 8.8×10^6^ ± 7.4×10^5^ and zero. **(D)** Hematoxylin and eosin (H&E) staining of testis cross-sections from 2-month old males. Left and middle panel size bars = 75 μm. Right panels are magnifications (2X) of the corresponding dashed boxes. **(E)** Elevated apoptosis in *Spo11^P306T^* testes. Testis cross-sections from 2-4 month old males were labeled with TUNEL and DAPI to detect apoptotic cells. Size bar = 75 μm. The number of seminiferous tubule cross sections with 5 or more TUNEL-positive cells is plotted below. Each data point corresponds to one testis. See Methods for details. **(F)** Quantification of primordial follicles from 3-week old ovaries. Each symbol represents one ovary. Averages: +/+ n=3, 1790 ± 38; *Spo11^P306T/^+* n=3, 1620 ± 122; *Spo11^P306T/P306T^*n=3, 794 ± 35; *Spo11^P306T/−^* n=3, 93 ± 18. All statistics were done using paired Student T-test. Average ± SEM.

Gross histology of *Spo11^P306T/P306T^* ovaries was unremarkable, but the overall oocyte reserve (primordial follicles) in 3 week old females was decreased by 56% (Fig. 2f). We surmise that although fecundity of mutant females did not decline over the course of the experimental time frame (<8 months), the oocyte reserve might become exhausted sooner in mutants than WT.

### Spo11^P306T/P306T^ spermatocytes have poor sex chromosome synapsis

SPO11-induced DSBs are critical for initiating homologous recombination repair that drives synapsis between homologous chromosomes. If synapsis fails profoundly, as in the case of *Spo11* nulls, spermatocytes will arrest with chromosomes in a zygotene-like configuration, resulting in complete sterility (Romanienko and Camerini-Otero 2000; Baudat et al. 2000b; Metzler-Guillemain and de Massy 2000). Since *Spo11^P306T/P306T^* males are fertile but exhibit dying spermatocytes and fewer sperm, this suggested that some fraction of spermatocytes might have abnormal DSB formation and thus some degree of defective synapsis. To test this, we immunolabeled spermatocyte chromosome surface spreads for the SYCP3 and SYCP1, which are proteins of the lateral and transverse elements of the synaptonemal complex (SC), respectively, and HORMAD2, a protein that is bound to unsynapsed SC axes but then is displaced upon synapsis (Wojtasz et al. 2009). In most *Spo11^P306T/P306T^* spermatocytes, the autosomes were synapsed properly by early pachynema; however, the sex chromosomes were unsynapsed in 33.3% (51/153, n=3 mice) of pachytene-diplotene cells, compared to 9.2% in WT (13/142, n=3) (Fig. 3a), suggesting a lack of DSB-induced PAR crossovers.

**Figure 3.**
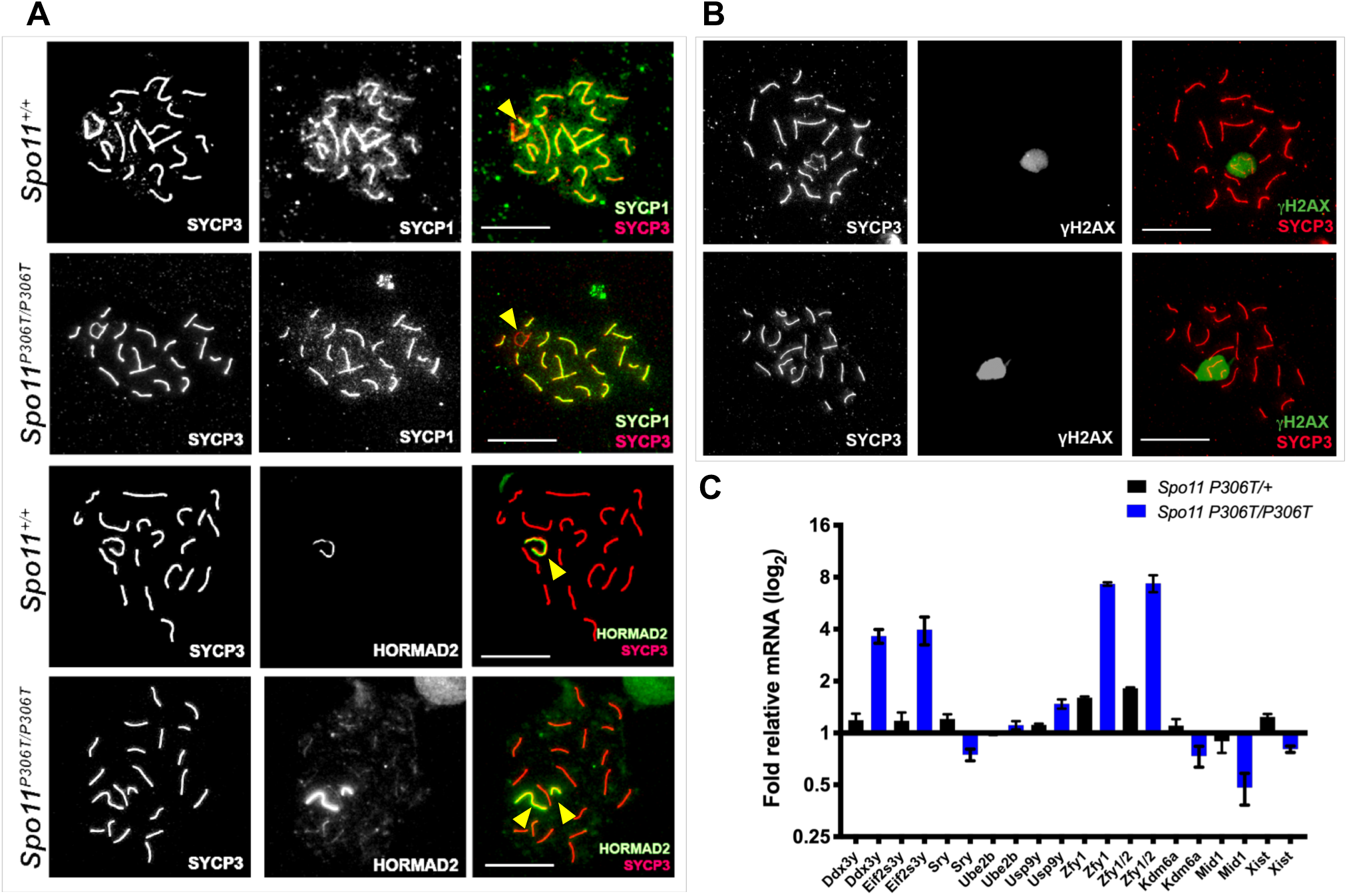
Sex chromosome pairing and silencing defects in *Spo11^P306T/P306T^* spermatocytes. **(A)** Representative pachytene chromosome surface spreads immunolabeled for indicated proteins. The 3^rd^ column is a merge of the first two columns. HORMAD2 labels unsynapsed regions of chromosomes, including the non-pseudoautosomal region of the XY pair. In the examples shown here, the X and Y chromosomes (yellow arrowheads) are not synapsed in the *Spo11^P306T/P306T^* spermatocytes (unstained in the PAR by the synapsis marker SYCP1; and entirely unpaired in the HORMAD2 example). Size bar = 20 μm. **(B)** Assessment of DNA damage and silencing in pachytene spermatocytes. Meiotic chromosome spreads were immunolabeled for γH2AX and SYCP3. Size bar = 20 μm. The mutant XY body shows normal staining for γH2AX, a marker of both DNA damage and heterochromatin, even in cases where there is XY asynapsis. For each genotype, 50 cells from each of 3 animals were examined. **(C)** RT-qPCR of X-linked genes from P15 testes relative to WT GAPDH. N=3 animals for each genotype.

Proper pairing and synapsis between the sex chromosomes leads to their heterochromatinization and transcriptional silencing, called meiotic sex chromosome inactivation (MSCI). Cytologically, the silenced XY is located at the periphery of the nucleus in a distinct structure called the XY body. Disruption of proper XY body formation, precipitated by failed synapsis at the PAR, can disrupt MSCI and lead to the expression of Y-linked genes (*Zfy1/2*) that lead to spermatocyte death (Royo et al. 2010). Because any meiotic defects that alter DSB repair or synapsis impact the silencing machinery needed for MSCI, the expression of these Y genes serve as a quality control mechanism to eliminate defective spermatocytes. Nevertheless, the XY body in mutant pachytene spermatocytes appeared intact as judged by immunolabeling of chromosome spreads with γH2AX (Fig. 3b), which intensely localizes to the XY body, and is essential for MSCI (Fernandez-Capetillo et al. 2003). Although the XY heterochromatinization appeared normal, we performed real-time quantitative reverse transcription-PCR (qRT-PCR) of select sex-linked genes to test for potentially aberrant expression in P15 testes, at developmental time at which the testes are enriched for pachytene spermatocytes produced during the first wave of spermatogenesis (Bellve et al. 1977). Although several genes were not differentially expressed between mutant and WT, we observed an ~8-fold higher level of the *Zfy1* and *Zfy2* spermatocyte “death genes” in *Spo11^P306T/P306T^* relative to WT (Fig. 3e) (Royo et al. 2010). At least two other genes were also expressed more highly in the mutant (*Ddx3y* and *Eif2s3y*). Taken together, the results thus far indicate that *spo11^P306T/P306T^* spermatocytes are capable of producing sufficient DSBs in a timely manner for autosome synapsis, but not always XY synapsis, and the latter may subtly disrupt MSCI. However, killing of spermatocytes by aberrant *Zfy1/2* expression typically occurs in mid-pachynema, not metaphase as indicated histologically (Fig. 2d). Thus, the *Spo11* mutation is likely causing more complex and subtle defects.

### Spo11^P306T/P306T^ meiocytes have altered timing of DSB formation and/or repair

To determine if SPO11^P306T^ impacts the number or processing of meiotic DSBs, we immunolabeled spermatocyte surface spreads with the ssDNA binding protein RPA2, and the RecA homologs RAD51and DMC1. There were no significant differences in RPA2 foci from the leptotene through late pachytene stages, although there were slightly fewer foci in mutant zygotene spermatocytes (87% of WT; Fig. 4a). RAD51 and DMC1 foci were also slightly (but significantly) decreased in mutant zygotene spermatocytes (Fig.4c-e), but were markedly increased in early pachynema, and especially prominently so on the sex chromosomes. By late pachynema, the number of DSBs was low in mutants, probably reflecting either rapid repair of the DSBs in early pachynema, or elimination of those spermatocytes by the pachytene checkpoint (see below). To determine if alterations to DSB number or timing were also altered in females, oocytes in late zygonema and beyond (obtained from newborns) were immunolabeled with RAD51, revealing similar trends as in spermatocytes (Fig. 5a,b). Late zygotene mutant oocytes had fewer RAD51 foci, but pachytene oocytes had slightly more.

**Figure 4.**
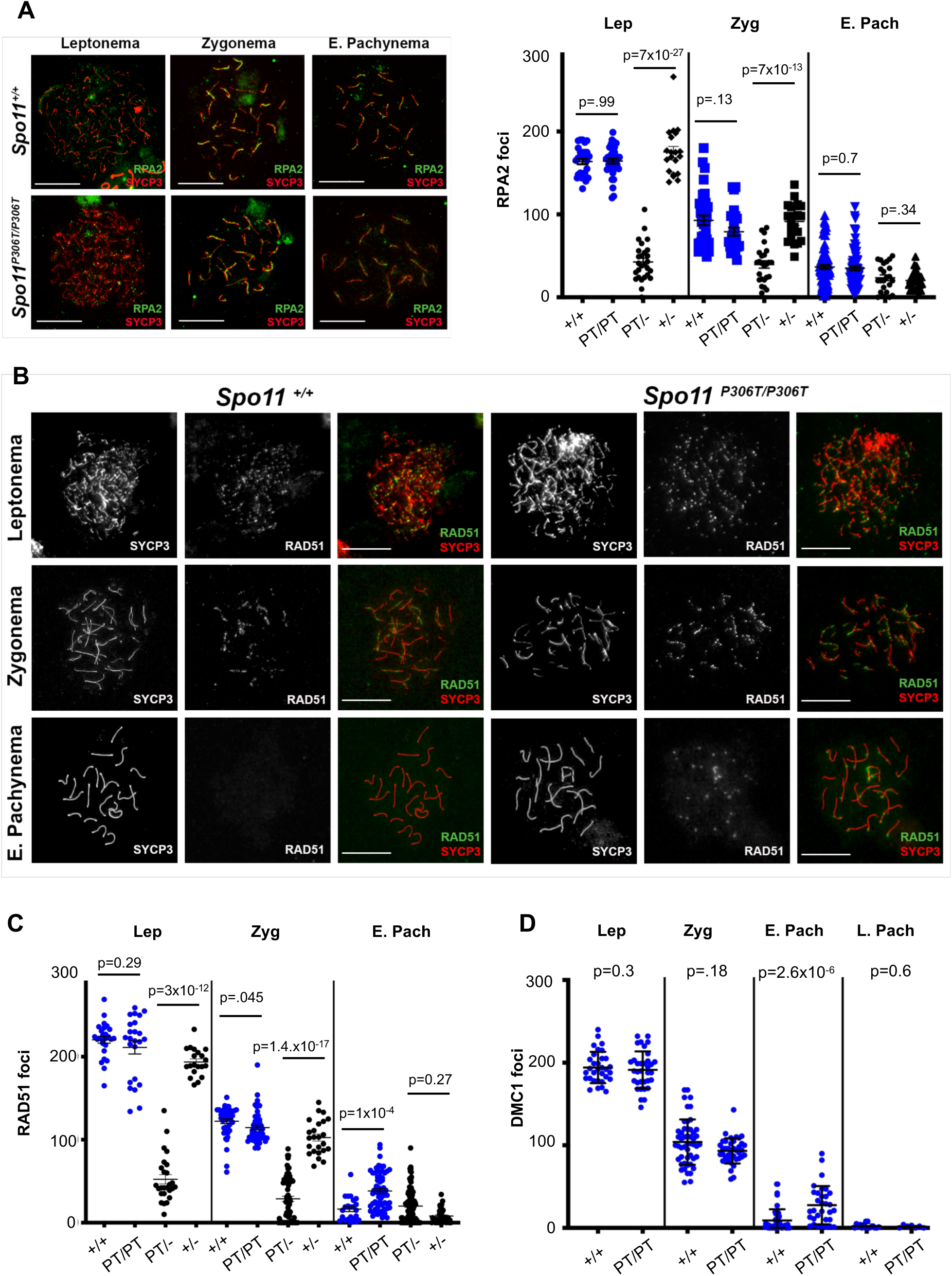
Assessment of DSB formation and repair in *Spo11^P306T/P306T^* mutants throughout meiotic prophase I. **(A)** Meiotic chromosome surface spreads immunolabeled with RPA2 and SYCP3 at different Prophase I substages. E, early. Here and in panel “b,” pachytene substage (early vs late) was determined on the basis of H1t staining (only late pachynema is positive for H1t; staining not included in these images). Size bar = 20μm. Quantification of RPA2 foci throughout Prophase I is plotted on the right. Actual numbers are as follows. Leptonema: +/+ (n=3 mice, cells=22, avg=164 ± 3.8), *Spo11^P306T/P306T^*(n=3, cells=27, avg=164 ± 3.5), *Spo11^P306T/−^* (n=2, cells=23, avg=39 ± 6.2), *Spo11^+/−^* (n=2, cells=20, avg=175 ± 6.6); Zygonema: +/+ (n=3, cells=36, avg=93 ± 5.9), *Spo11^P306T/P306T^*(n=3, cells=27, average=81 ± 3.8), *Spo11^P306T/−^* (n=2, cells=28, avg=37 ± 4.4), *Spo11^+/−^* (n=2, cells=29, avg=92 ± 4.7); Early pachynema: +/+ (n=3, cells=87, avg=37 ± 2.4), *Spo11^P306T/P306T^* (n=3, cells=72, avg=35 ± 2.7), *Spo11^P306T/−^* (n=2, cells=29, avg=24 ± 4.5), *Spo11^+/−^* (n=2, cells=24, avg=20 ± 2.2). **(B)** Meiotic chromosome spreads immunolabeled for RAD51 foci during indicated prophase substages. Size bar = 20μm. **(C)** Quantification of RAD51 foci at prophase I substages. Actual numbers are as follows: Leptonema: +/+ (n=4, cells=26, avg=220 ± 4.1), *Spo11^P306T/P306T^*(n=5, cells=24, avg=211 ± 7.8), *Spo11^P306T/−^(n=5*, cells=23, avg=70 ± 7.9), *Spo11^+/−^* (n=2, cells=17, avg=193 ± 4.3); Zygonema: +/+ (n=4, cells=43, avg=122 ± 2.8), *Spo11^P306T/P306T^*(n=5, cells=52, avg=115 ± 2.6), *Spo11^P306T/−^* (n=5, cells=59, avg=30 ± 3.6), *Spo11^+/−^* (n=2, cells=25, avg=101 ± 4.7); Early pachynema: +/+ (n=4, cells=23, avg=14.2 ± 2.6), *Spo11^P306T/P306T^*(n=5, cells=70, avg 40.8 ± 3.4), *Spo11^P306T/−^* (n=5, cells=80, avg= 15 ± 1.2), *Spo11^+/−^* (n=2, cells=37, avg=8.7 ± 1.5). **(D)** Quantification of DMC1 levels at prophase I substages. Leptonema: WT (n=3, cells=32, avg=194 ± 3.3) vs mutant (n=3, cells=32, avg=189 ± 3.6); Zygonema: WT (n=3, cells=33, avg=99 ± 4.6) vs mutant (n=3, n=33, avg=92 ± 2.8); Early pachynema: WT (n=3, cells=47, avg=8.9 ± 2.0) vs mutant (n=3, cells=32, avg=27 ± 4.1, p=2.6×10^−5^); Late pachynema: WT (n=3, cells=20, avg=2.2 ± 0.89) vs mutant (n=3, cells=20, avg=1.6 ± 0.51). All statistics were done using paired Student T-test. Avg ± SEM.

**Figure 5.**
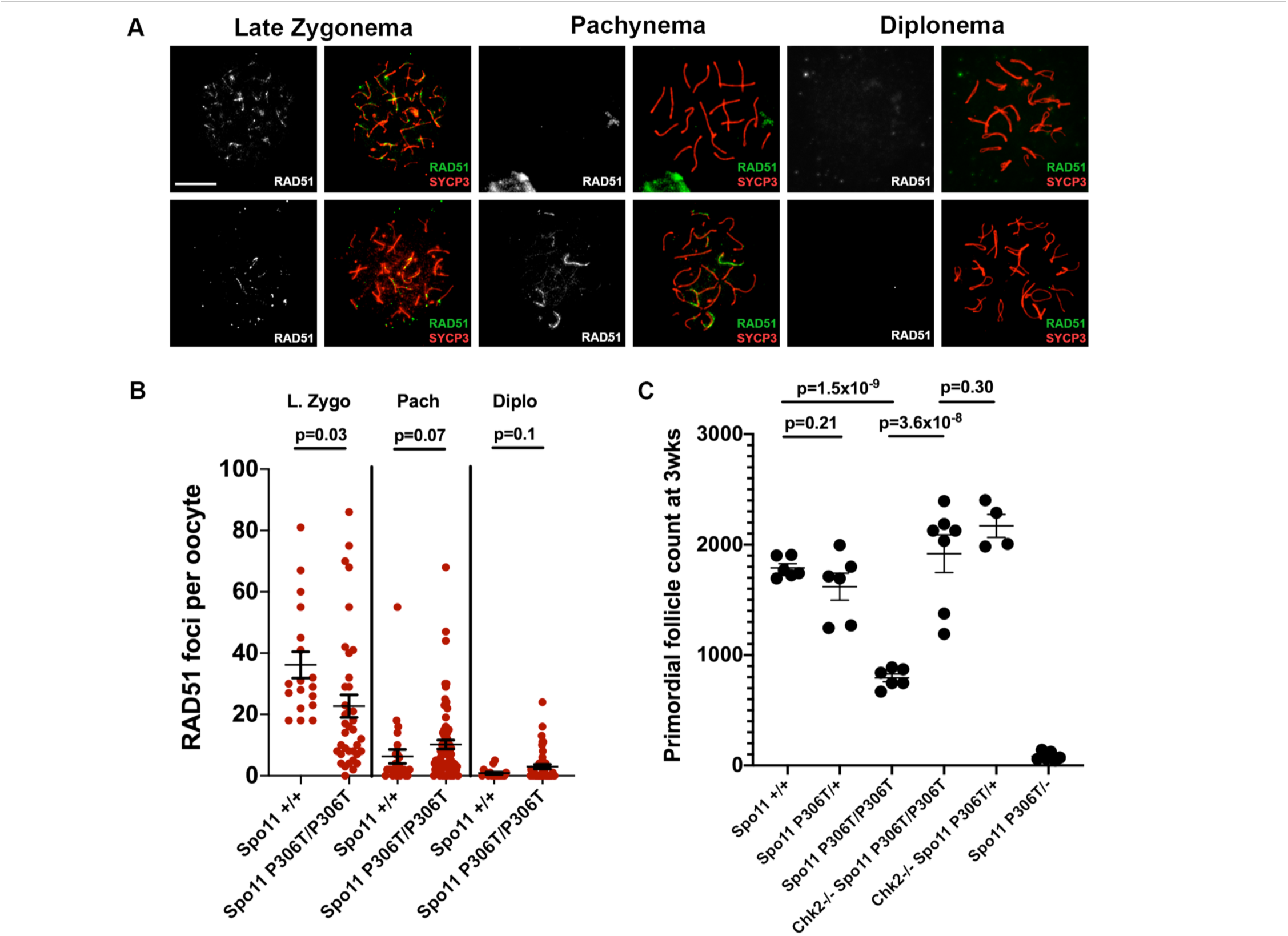
Oocytes have delayed DSB formation but die from unrepaired DSBs in late Prophase I. **(A)** Representative P0 oocyte chromosome spreads immunolabeled with RAD51 and SYCP3. Size bar = 25μm. **(B)** Quantification of RAD51 foci. Late zygonema: WT (n=2, cells=18, avg=36 ± 4.3) vs mutant (n=2, cells=37, avg=23 ± 3.7); pachynema: WT (n=2, cells=24, avg= 5.5 ± 1.1) vs mutant (w=2, cells=70, avg=10 ± 1.5); diplonema: WT (n=2, cells=20, avg=0.9 ± 0.3) vs mutant (n=2, cells=48, avg=3 ± 0.7). Avg ± SEM. **(C)** Quantification of primordial follicles in 3 week old females, with or without an intact DNA damage checkpoint. The leftmost 3 datasets and the Rightmost dataset are identical to those in Fig. 2F. Each data point is one ovary, and both ovaries were counted from each female. Statistics done by Student’s T-test.

These findings suggest that although SPO11^P306T^ protein and or activity is reduced in leptonema and zygonema, it is sufficient to make enough DSBs of a nature that catalyze recombination-driven pairing/synapsis. However, the basis for abnormal presence of RAD51/DMC1 foci into late pachynema in mutants is uncertain. The following are possible explanations: 1) the DSBs formed are of a nature that makes them more difficult to repair, or shunt them into a later-acting DNA repair pathway; 2) DSB production is not normally downregulated as prophase I progresses; and/or 3) the RAD51/DMC1 foci do not actually reflect DSBs.

To test the latter possibility, and also address whether the persistence of apparent DSBs into late pachynema contributes to observed gamete loss, we generated *Spo11^P306T/P306T^ Chek2^−/−^* mice and quantified sperm and the ovarian reserve. *Chek2* is a key element of the meiotic DNA damage checkpoint pathway that eliminates oocytes bearing ~10 or more DSBs (RAD51 foci) in late pachynema (Rinaldi et al. 2017; Bolcun-Filas et al. 2014), a level exceeded in 28% of pachytene oocytes (Fig. 5b). *Chek2* ablation dramatically rescued primordial follicles in three-week ovaries (Fig. 5C). In contrast, *Chek2^−/−^ Spo11^P306T/P306T^* males remained subfertile, with unchanged sperm levels (data not shown). However, this is unsurprising, because disruption of MSCI is the major cause of spermatocyte elimination, essentially acting upstream (temporally) of the DNA damage checkpoint (Pacheco et al. 2015).

The average number of RAD51 foci in mutant diplotene oocytes decreased from pachytene levels, suggest that either the pachytene oocytes with a DSB burden above the threshold for checkpoint activation were eliminated before progression to diplonema, or that DSBs were repaired in this interval. In conclusion, the excess RAD51 foci in pachytene cells appear to correspond with unrepaired DSBs, although the data presented thus far did not allow us to distinguish between the possibilities 1 and 2 listed above regarding the timing and repairability of the DSBs in *Spo11^P306T/P306T^* mice.

### The Spo11^P306T^ allele decreases Class I crossover formation

In normal mice, ~10% of SPO11-catalyzed DSBs are repaired as crossovers (COs). The resulting chiasmata are essential for tethering homologous chromosomes at the meiotic prophase I metaphase plate, ensuring segregation of homologs to opposite daughter cells (Gray and Cohen 2016). As mentioned earlier, testis histology revealed the presence of abnormal metaphase I spermatocytes that appeared to have disorganized chromatin or lagging chromosomes at the metaphase plates (Fig. 2d), suggesting a possible defect in crossing over. There are two known CO mechanisms in mammals, the major pathway (“Class I”), accounting for ~90% of all COs, being mediated by the *MutL* homologs MLH1 and MLH3 (Guillon et al. 2005; Rogacheva et al. 2014).

To test whether mutants had defects in crossing over, we quantified the number of MLH1 foci in spermatocytes. *Spo11^P306T/P306T^* spermatocytes averaged significantly fewer MLH1 foci than WT (19.6±0.2 vs 23.7±0.1, respectively) (Fig. 6a,b). Strikingly, mutants exhibited an increase in chromosomes without any foci. Whereas WT pachytene spermatocytes rarely a chromosome pair lacking an MLH1 focus (avg. of 0.62 ± 0.08), mutants averaged 3.9 ± 0.3. This failure to distribute an “obligate” crossover to every chromosome may not only cause aneuploid daughter cells, but also would be predicted to activate the spindle checkpoint in some spermatocytes to trigger their elimination.

**Figure 6.**
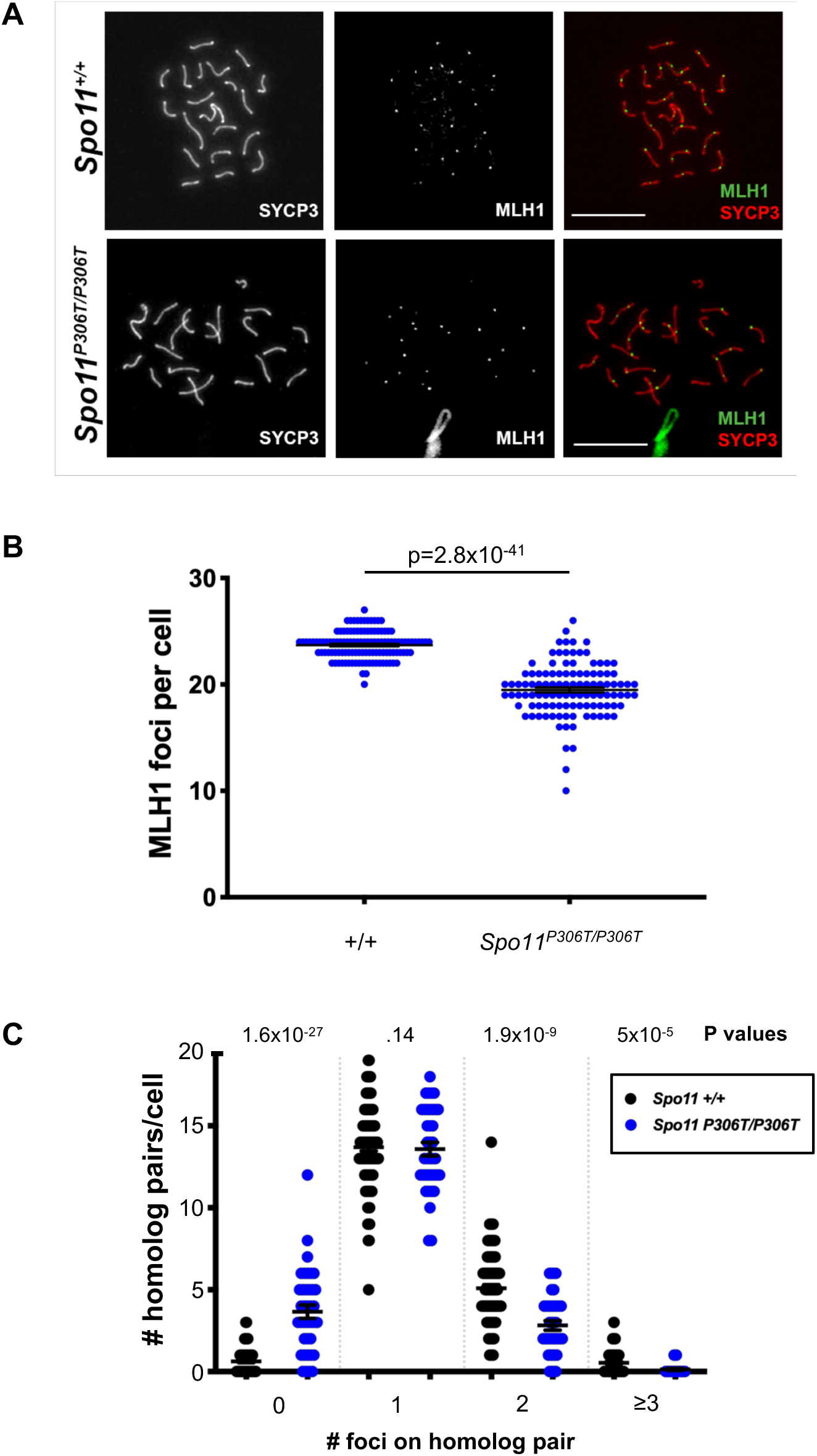
*Spo11^P306T/P306T^* spermatocytes have fewer MLH1 foci, indicative of reduced crossovers. **(A)** Chromosome spreads of pachytene spermatocytes immunolabeled with MLH1 and SYCP3. Size bar= 20μM. **(B)** Quantification of MLH1 foci. Actual numbers are as follows: WT (n=3, cells=118, avg=23.7 ± 0.12), and *Spo11^P306T/P306T^*(n=4, cells=108, avg=19.6 ± 0.2). Avg ± SEM. **(C)** *Spo11^P306T/P306T^* spermatocytes have an increased incidence of chromosomes lacking an apparent crossover (MLH1 focus). N=3 mice for each genotype. For WT and mutant, 102 and 58 cells were scored, respectively. All statistics were done using paired Student T-test. Avg ± SEM.

### Genetic evidence that Spo11^P306T^ is a hypomorphic allele

The defects in *Spo11^P306T/P306T^* mice presented thus far are enigmatic. On one hand, one might expect SPO11 defects to cause a decrease in DSBs, leading to autosomal asynapsis from reduced recombination. There was some evidence for this in the data showing modestly decreased RAD51/DMC1 foci in early prophase I, and reduced crossovers. On the other hand, we observed elevated DSBs in pachytene spermatocytes, raising the possibility that it has increased or dysregulated activity. We reasoned that if *Spo11^P306T^* were a hypermorph, then removing a dose would ameliorate the phenotypes, by decreasing DSBs. Conversely, if it were a hypomorph, then removing a dose would exacerbate the phenotypes. Accordingly, we bred and analyzed mice bearing *Spo11^P306T^* in *trans* to a *Spo11* null allele.

Both male and female *Spo11^P306T/−^* mice had more severe phenotypes than *Spo11^P306T^* homozygotes. Males exhibited smaller testes (Fig. 2b), no epididymal sperm (Fig. 2c), and seminiferous testes devoid of postmeiotic cells (Fig.S1a). Females had severely depleted oocyte reserves (Fig. 2f). Consistent with the severe histopathology, matings of *Spo11^P306T/−^* males or females to WT mates were non-productive. At this level of characterization, *Spo11^P306T/−^* mice resemble *Spo11* nulls that exhibit infertility in both sexes associated with meiotic arrest (Baudat et al. 2000a; Romanienko and Camerini-Otero 2000), thus suggesting that *Spo11^P306T^* is hypomorphic. To test this, we immunolabeled *Spo11^P306T/−^* and *Spo11^+/−^* chromosome spreads for DSB markers. RPA foci in *Spo11^P306T/−^* leptotene and zygotene spermatocytes had only 78% and 60% the numbers of foci, respectively, compared to *Spo11^+/−^* spermatocytes (Figs. 2a, S1b). Similar results were obtained with RAD51, where *Spo11^P306T/−^* spermatocytes averaged 36% and 30% the number of foci present in *Spo11^+/−^* leptotene and zygotene spermatocytes, respectively (Figs. 4c, S1c).

Despite the decreased number of DSBs, it was evident that *Spo11^P306T/−^* spermatocytes had substantial (though incomplete) synapsis of homologous chromosomes. *Spo11^P306T/−^* spermatocytes immunolabeled with HORMAD2, which decorates the length of unsynapsed SC axial elements and is removed upon synapsis (Wojtasz et al. 2009), revealed defects ranging from pachytene cells with 3+ partially synapsed homologous chromosomes to diplotene cells with complete synapsis (Fig. S1d). Among spermatocytes with asynapsed chromosomes or chromosome subregions, sex chromosome were always affected.

### Evidence that the mutation affects timely DSB formation

Having provided evidence that germ cell loss in mutants was due to presence of DSBs in pachynema that triggered the DNA damage checkpoint, we addressed the hypotheses (articulated above) regarding the mechanism leading to a surfeit of late DSBs, i.e., they are more difficult to repair *vs* they are made later than normal. To gain insight into these possibilities, we examined patterns of SPO11 accessory protein localization on spermatocyte chromosomes throughout prophase I.

MEI4 is a conserved protein that is required for DSB formation. It localizes to foci along asynapsed chromosome axes in a SPO11-independent manner, and is apparently displaced upon initiation of DSB repair (RAD51/DMC1 loading) and local synapsis (Kumar et al. 2010; Kumar et al. 2015). In contrast to RAD51/DMC1 foci which were decreased compared to WT, MEI4 foci in *Spo11^P306T/P306T^* zygotene spermatocytes were modestly increased (Fig. 7a,b). In *Spo11^P306T/−^* spermatocytes, MEI4 foci were elevated in both leptonema and zygonema. Additionally, MEI4 persisted along large stretches of chromosomes in early pachynema (Fig. 7a,b). This is suggestive of delayed DSB formation in mutants. MEI4 removal has been suggested to negatively feed back on DSB production (Kumar et al. 2015). If true, MEI4’s elevated numbers during later prophase I stages may reflect delayed DSB formation and resulting in lack of feedback inhibition.

**Figure 7.**
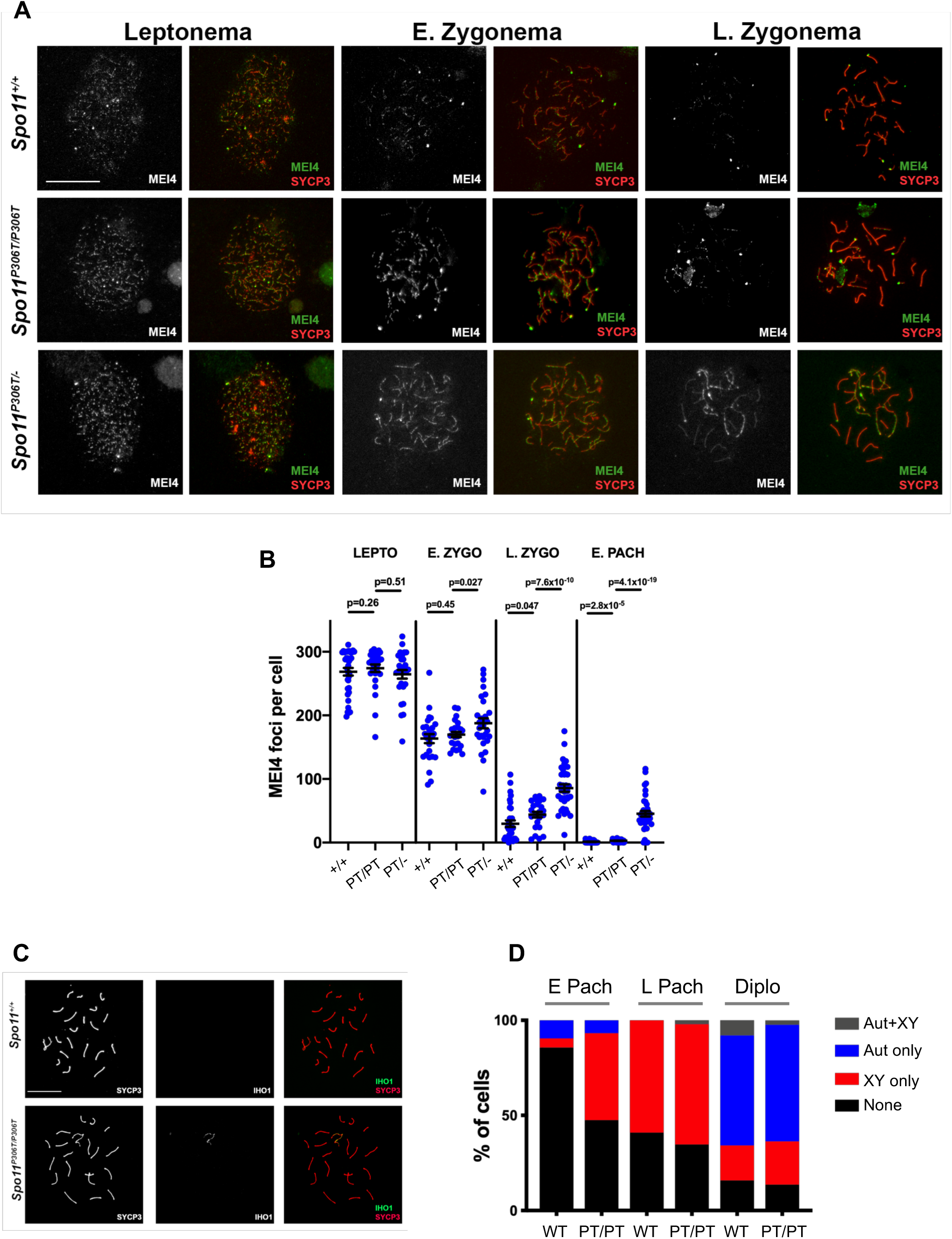
Pre-DSB recombinosomes persist on chromosome axes of *Spo11^P306T/P306T^*and *Spo11^P306T/−^*. **(A)** Chromosome surface spreads of spermatocytes immunolabeled with MEI4 and SYCP3. Size bar = 20μm. **(B)** Quantification of MEI4 foci at different phases of prophase I. Actual numbers are as follows. Leptonema: WT (n=2 mice, cells=27, avg=163 ± 7.1), *Spo11^P306T/P306T^* (n=3, cells=25, avg=169 ± 4.1), and *Spo11^P306T/−^* (n=2, cells=29, avg=188 ± 7.9); Early Zygonema: WT (n=2, cells=32, avg=29.7 ± 5.1), *Spo11^P306T/P306T^*(n=3, cells=25, avg=44 ± 4.3) and *Spo11^P306T/−^* (n=2, cells=35, avg=86 ± 5.9); Late Zygonema: WT (n=2, cells=51, avg=1.2 ± 0.26), *Spo11^P306T/P306T^* (n=3, cells=48, avg=2.8 ± 5.9) and *Spo11^P306T/−^* (n=2, cells=38, avg=45 ± 4.5). Pachynema: WT (n=2, cells=51, avg= 1.2 ± 0.3), *Spo11^P306T/P306T^* (n=3, cells=48, avg=2.8 ± 5.9), and *Spo11^P306T/−^* (n=2, cells=38, avg=45.4 ± 4.5). **(C)** Chromosome spreads of pachytene spermatocytes immunolabeled with IHO1 and SYCP3. Size bar = 20μm. **(D)** Percentage of early- and late pachytene and diplotene spermatocytes with indicated IHO1 localization patterns. Early pachynema: WT (n=2, cells=40) and *Spo11^P306T/P306T^* (n=3, cells=60); late pachynema: WT (n=2, cells= 30) and *Spo11^P306T/P306T^* (n=3, cells=60); diplonema: WT (n=2, cells=30) and *Spo11^P306T/P306T^* (n=3, cells=30). Aut = autosomes; PT = *Spo11^P306T/P306T^*. All statistics were done using paired Student T-test. Avg ± SEM.

Interestingly, we also observed abnormal persistence of IHO1 (interactor of HORMAD1) foci in the *Spo11^P306T/P306T^* spermatocytes into early pachynema, but only on the sex chromosomes (Fig. 7c,d). IHO1 is required for DSB formation, and complexes with HORMAD1, MEI4, and another SPO11 accessory protein, REC114 (Stanzione et al. 2016). Normally, IHO1 is removed from most synapsed regions of autosomes, and also from both synapsed and unsynapsed regions of the sex chromosomes upon pachytene entry (Stanzione et al. 2016), consistent with our observations in WT controls (Fig. 7c,d). DSB formation on the sex chromosomes is dependent upon progress of autosomal DSB formation and repair; DSBs only appear at the PAR after ~70% of RAD51 foci are displaced from autosome axes (Kauppi et al. 2011). Our observations of elevated XY asynapsis, and late XY DSBs and IHO1 foci on the XY are consistent with an overall delay in the progress of DSB formation and repair in the mutant, ultimately leading to the elimination of some spermatocytes.

## DISCUSSION

A major challenge in human genomics is to elucidate the effects of variants of unknown significance (VUS). Presumably, it should be simplest to predict the effects of those that alter protein sequence (e.g nonsynonymous SNPs, or nsSNPs). Numerous algorithms have been developed to do just that, but experimental testing of putative deleterious nsSNPs has called their effectiveness into question (Miosge et al. 2015; Singh and Schimenti 2015; Wang et al. 2018). Nevertheless, it must be recognized that these algorithms do not predict phenotype; rather, they predict the likelihood that an amino acid change impacts the protein in a deleterious manner. Of course, the biochemical effects of a deleterious mutation upon the protein could range from complete or partial loss-of-function, to hyperactivity or gain-of-function, and in most cases this is impossible to predict. More difficult to predict is the *in vivo* phenotype of an organism bearing a VUS, especially in the absence of genetic data.

This study of the *Spo11^P306T^* allele is part of a larger-scale project to identify human infertility alleles, using a combination of *in silico* predictions and *in vivo* modeling in mice. This allele models a SNP that was selected by virtue of the following criteria: 1) It resides in a gene known to cause infertility when knocked out in mice, and is an nsSNP; 2) It is a bona fide segregating SNP in the population as judged by its presence in 15 sequenced people and having passed gnomAD browser filters; 3) its frequency is not high (which would be inconsistent with it being an infertility allele); and 4) it was scored by multiple algorithms as being damaging to the SPO11 protein. Based on the last two criteria alone, in the absence of any other genetic information, it would be tempting to conclude that an infertile proband bearing this mutation, either homozygously or in *trans* to another predicted deleterious allele, would be causative for fertility defects. *A priori*, it would be unlikely for dominant infertility alleles to persist in a population, although there is evidence for such alleles to affect only one sex (Bannister et al. 2007; Singh and Schimenti 2015).

Our finding that mutant homozygotes are fertile would, in isolation, lead one to conclude that *Spo11^P306T^* is not an infertility allele, and that the *in silico* predictions were incorrect. However, closer analysis revealed that *Spo11^P306T^* not only caused sterility in *trans* to a null allele, but also that homozygotes had a lower sperm counts and a reduced ovarian reserve. As no human homozygotes have been described (to our knowledge), it is unclear whether the effects would be similar. Nevertheless, it is possible that the allele could contribute to premature ovarian failure if a woman’s oocyte reserve was similarly reduced (~50%). Coupled with other genetic or environmental defects, men with the mutation may have decreased sperm counts that could cause fertility problems. Even if fertility was not substantially impacted in people, our observations of defects in recombination, most notably a reduction in crossing-over to a level below the number of chromosomes, raises the possibility that the allele could contribute to chromosome imbalances in offspring that typically lead to pregnancy loss. In sum, our work underscores the importance of performing the detailed phenotyping to fully understand the consequences of a suspected deleterious VUS. Unfortunately, in the case of gametogenesis, such phenotyping is not simple, and casts doubt upon whether alternatives such as *in vitro* gametogenesis will ever recapitulate the *in vivo* situation sufficiently effectively to detect such nuanced phenotypes.

The proper generation and repair of SPO11-dependent DSBs is essential for successful pairing and segregation of meiotic chromosomes. Multiple aspects of the process are finely regulated to enable successful outcomes. These include generating a sufficient number of DSBs (normally ~200-250 in mouse spermatocytes) to drive homologous recombination (HR)-driven chromosome pairing, predominantly (~90%) via noncrossover (NCO) recombination, and reserving ~10% to be repaired as crossovers (COs) such that each chromosome receives at least one. As discussed above, the latter is crucial because chromosome pairs that aren’t tethered at the metaphase I plate by a CO can segregate to the same daughter cells, potentially leading to aneuploid embryos. Laboratory mice, which have 20 chromosomes, have ~26 COs per meiosis (Gray and Cohen 2016) which are distributed nonrandomly across all chromosomes, such that each chromosome gets at least 1 CO. This “crossover assurance” is attained via interference (Broman et al. 2002) and a potentially related phenomenon known as “crossover homeostasis” in which CO numbers are maintained at the expense of NCOs when SPO11 DSBs are reduced genetically (Martini et al. 2006).

The reduction of COs in *Spo11^P306T/P306T^* meiocytes suggests that CO homeostasis was disrupted. This may be a consequence of either a reduction of DSBs below a threshold that can be compensated by the homeostasis mechanism, or that activation of the mechanism is delayed along with DSB induction in the mutant. We tend to favor the latter, since at no point were the level of DSBs, as measured by RAD51 foci, lower in *Spo11^P306T/P306T^* spermatocytes than in *Spo11^+/−^*, the latter of which does not suffer from loss of COs (Carofiglio et al. 2013). Rather, our evidence of slightly reduced (relative to WT) DSBs in leptonema and zygonema, but markedly increased numbers in pachynema, suggested that the *Spo11^P306T^* mutation causes delayed DSB formation. The delay is likely not a consequence of reduced levels of SPO11^P306T^ protein (Fig. S2), but possibly reduced catalytic activity or decreased ability to interact with accessory proteins. The proline 306 amino acid is conserved from yeast to mammals, residing in the catalytic TOPRIM domain of related topoisomerase and topoisomerase-like proteins (Robert et al. 2016).

We suggest that the presence of excess DSBs in early pachynema in the mutant likely reflects abnormally late induction of DSBs. The alternative that there is defective repair seems unlikely, since SPO11 is not involved in DSB repair, and is removed from DSB sites to be replaced by recombination proteins (including RAD51 and DMC1) that conduct the repair and mark the late DSBs. Multiple lines of evidence indicate that regulation of DSB formation involves a negative feedback loop mediated by the ATM kinase. Mouse spermatocytes lacking ATM have high levels of RAD51 foci and chromosome fragmentation, leading to meiotic prophase I arrest (Xu et al. 1996). This arrest could be bypassed by *Spo11* heterozygosity (Bellani et al. 2005), and, in conjunction with the observation that *Atm^−/−^* mice have increased levels of SPO11-oligonucleotide complexes, led to the conclusion that ATM senses DSB levels and downregulates SPO11 activity with appropriate timing in prophase I (Lange et al. 2011). The activity of such a mechanism could explain, for example, why *Spo11^+/−^* spermatocytes have at least 70% of WT DSB levels, rather than 50% (Lange et al. 2011; Cole et al. 2012; Kauppi et al. 2013; Neale et al. 2005). The exact mechanism of this regulation is unknown, but may involve either direct or indirect regulation of SPO11 activity. For example, ATM may modulate accessory protein levels or activity as prophase I progresses (Cole et al. 2012; Lange et al. 2011). We hypothesize that the abnormal presence of late DSBs in *Spo11^P306T/P306T^* spermatocytes is a consequence of delayed activation of this negative feedback mechanism, which occurs as a result of compromised SPO11 catalytic activity.

Analysis of *Spo11^P306T/−^* mice revealed some remarkable phenotypes. They exhibited only 20-40% normal levels of RPA2 and RAD51 foci in leptotene and zygotene spermatocytes, and the histological and fertility phenotypes were catastrophic compared to *Spo11^P306T/P306T^*. Nevertheless, there were substantial levels of synapsis.

Despite the greatly reduced apparent DSB formation as judged by these markers, *Spo11^P306T/−^* spermatocytes, like *Spo11^P306T/P306T^* spermatocytes, still had evidence of persistent or ongoing DSB formation in early pachynema. Presumably, this also reflects a delayed or absent negative feedback for DSB production, and/or lack of positive stimulation of DSB formation. Previous studies suggested that 40% of normal DSB activity is required for proper synapsis and meiotic progression (Faieta et al. 2016). Our data with *Spo11^P306T/−^* spermatocytes is in general agreement, considering only a subset of cells appear to survive through meiosis, which might represent those cells at the higher end of the DSB formation range.

Both human and mouse spermatocytes produce two SPO11 isoforms, alpha and beta, resulting from alternative splicing of exon 2 (Romanienko and Camerini-Otero 1999). *Spo11β* predominates in leptonema and zygonema, while only *Spo11α* transcripts are made in pachynema (Bellani et al. 2010)(Kauppi et al. 2011), and the latter is especially crucial for inducing at least one DSB into the psudoautosomal region (PAR) of the mouse XY pair (Kauppi et al. 2011). Interestingly, a phenotype similar to that in *Spo11^P306T/P306T^* males has been observed in a mice expressing only the beta isoform from a transgenic in a *Spo11^−/−^* background (Kauppi et al. 2011; Kauppi et al. 2013). These males have WT levels of autosomal DSBs and normal autosomal synapsis and COs, but ~70% of spermatocytes have asynapsed sex chromosomes due to failure to induce DSBs. We conjecture that the delay in autosomal SPO11 induction, presumably by decreased enzymatic activity of SPO11β, either delays expression of SPO11 *α* and/or that the enzymatic activity of the alpha isoform is also decreased by virtue of containing the altered TOPRIM domain.

The *Spo11^P306T^* mutation also had dramatic effects on oogenesis. *Spo11^P306T/P306T^* females exhibit 50% oocyte loss at three-weeks of age, while *Spo11^P306T/−^* females have almost complete loss, with only ~5% of WT levels, rendering them infertile. They also showed a delay in DSB formation (elevated RAD51 foci in zygonema), but elevated DSBs in pachynema, suggesting that a feedback mechanism akin to that in spermatocytes also exists in oocytes. Our results showed that the delayed induction, and/or late repair of these DSBs, led to activation of the CHEK2-mediated DNA damage checkpoint (Bolcun-Filas et al. 2014). It is possible that the oocytes that avoid elimination by this checkpoint (by having a number of DSBs that is below the threshold of checkpoint detection) are more prone to Meiosis I chromosome segregation errors stemming from asynapsed chromosomes (Fig. 5). If so, this mutation may lead to increased pregnancy loss.

## Acknowledgements

This work was supported by a grant from the National Institutes of Health (R01 HD082568 to JCS), and contract CO29155 from the NY State Stem Cell Program (NYSTEM). The authors would like to thank R. Munroe and C. Abratte of Cornell’s transgenic facility for generating the *Spo11^P306T^* allele, Scott Keeney for providing SPO11 antibody, and Attila Toth for providing MEI4, IHO1, and HORMAD2 antibodies.

## MATERIALS AND METHODS

### Generation of *Spo11^P306T^* Mice by CRISPR/Cas9 Genome Editing

The DNA template for making sgRNA was generated using a cloning-free overlap PCR method, essentially as described (Varshney et al. 2015). The guide RNA sequence corresponding to Spo11 was as follows: GACCAAGCCATCTGATTGTT. The DNA template was reverse-transcribed into RNA using Ambion MEGAshortscript T7 Transcription Kit (cat#AM1354), then purified using Qiagen MinElute columns (cat#28004). For pronuclear injection, the sgRNA (50 ng/μL), ssODN (50 ng/μL, IDT Ultramer Service), and Cas9 mRNA (25 ng/μL, TriLink) were co-injected into zygotes (F1 hybrids between strains FVB/NJ and B6(Cg)-Tyr^c-2J^/J) then transferred into the oviducts of pseudopregnant females. Founders carrying at least one copy of the desired alteration were identified and backcrossed into FVB/NJ. Initial phenotyping was done after one backcross generation and additional phenotyping was done with mice backcrossed at least two or more generations.

### Mice and genotyping

All animal use was conducted under protocol (2004-0038) to J.C.S. and approved by Cornell University’s Institutional Animal Use and Care Committee. Other than the *Spo11^P306T^* allele reported here, mutant mice used in this study were *Atm^tm1Awb^* (abbreviated *Atm^−/−^*) (Barlow et al. 1996), *Chk2*^tm1Mak^(abbreviated *Chk2^−/−^*) (Hirao et al. 2002), and *Spo11^tm1Mjn^* (abbreviated *Spo11^−/−^*) (Baudat et al. 2000a).

Crude lysates for PCR were made from small tissue biopsies (tail, toe or ear punches) as described (Truett et al. 2000). Genotyping primers are listed in Supplementary Table 1. PCR reactions were as follows: initial denaturation at 95° for 5 min, then 30 cycles of 95° for 30 sec, 58° for 30 sec, 72° for 30 sec, and final elongation at 72° for 5 min. For identification of *Spo11^P306T^* mutants, amplicons were digested with restriction enzyme HaeIII (NEB) at 37°C for 2h and products were analyzed on high percentage agarose gels. The WT products yield 247bp + 62bp bands, and the *Spo11^P306T^* allele yields bands of 125bp + 122bp + and 62bp.

### Sperm Counts

Cauda epididymides (both sides) were collected from 8-week old males, minced in PBS, then incubated at 37°C for 10 minutes to allow sperm to swim out. Sperm solutions were diluted into 10 mL and counted on a hemocytometer.

### Histology and Primordial Follicle Quantification

Testes were collected from 8-week males, and ovaries were collected from 3-week females. They were fixed in Bouin’s for 24h, washed in 70% ethanol for 24h, then embedded in paraffin. Testes were sectioned at 6 μm and stained with hematoxylin and eosin (H&E). Ovaries were serial sectioned at 6 μm, stained with H&E, and primordial follicles were counted in every fifth section. Final follicle counts per ovary were calculated as previously described (Myers et al. 2004). Statistical analysis was done with a two-tailed Student’s *t*-test using Prism 7 software (Graphpad).

### TUNEL assay

Testes were collected from 8-week males and fixed using 4% paraformaldehyde, embedded in paraffin, and sectioned at 6 μm. The Click-IT Plus TUNEL Assay for In Situ Apoptosis Detection Alexa Fluor™ 488 dye kit (ThermoFisher Scientific, #C10617) was used. Quantification as shown in Fig. 2E was performed as follows. Between 3 and 5 random sections of each testis was TUNEL labeled, and the number of tubules per cross section (~200) with ≥5 TUNEL+ cells was counted. The overall percentage of such tubules in all cross those sections in a testis was then determined.

### Immunocytochemistry of Meiotic Chromosomes

We used a published protocol (McNairn et al. 2017). In brief, testes from 8-12 week old males were detunicated and minced in MEM media. Spermatocytes were hypotonically swollen in 4.5% sucrose solution and lysed in 0.1% Triton X-100/0.02% SDS/2% formalin. Slides were washed and immediately stained or stored at −80°C. The blocking buffer used was 5% goat serum diluted in PBS/0.1% Tween20 and slides were blocked for 1h at room temperature. Primary antibodies were incubated for overnight at 4°C and dilutions used were anti-SYCP3 (1:600, Abcam, #ab15093), anti-SYCP3 (1:600, Abcam, #ab97672), anti-SYCP1 (1:400, Abcam, #ab15090), anti-RPA2 (1:100, gift from J.W.), anti-DMC1 (1:100, Abcam, #ab11054), anti-RAD51 (1:100, Millipore Sigma, #PC130-100UL), anti-HORMAD2 (gift from A.T.), anti-MEI4 (1:200, gift from A.T.), anti-IHO1 (1:200, gift from A.T.), anti-MLH1 (1:100, BD Pharmingen, #554073), anti-RNA Polymerase II (1:500, Millipore Sigma, #05-623), and anti-phospho-H2A.X (1:1000, Millipore, #16-193). Secondary antibodies were incubated at room temperature for 1h. Secondary antibodies used were goat anti-mouse IgG Alexa Fluor 488 (1:1000, ThermoFisher Scientific, A-11001), goat anti-mouse IgG Alexa Fluor 594 (1:750, ThermoFisher Scientific, A-11032), goat anti-rabbit IgG Alexa Fluor 488 (1:1000, ThermoFisher Scientific, #R37116), goat anti-rabbit IgG Alexa Fluor 594 (1:750, ThermoFisher Scientific, A-11012), goat anti-guinea pig IgG Alexa Fluor 594 (1:750, ThermoFisher Scientific, A-11076), and goat anti-guinea pig IgG Alexa Fluor 647 (1:1000, ThermoFisher Scientific, A-21450). Images were acquired with an Olympus microscope using cellSens software (Olympus).

Foci were quantified using ImageJ with plugins Cell Counter (Kurt De Vos) and Nucleus Counter. All data was analyzed statistically using Prism 7 (GraphPad).

### Real-time Quantitative PCR

Total RNA was isolated from tissue using the E.Z.N.A. kit (Omega Biotek) according to manufacturer’s protocol. A total of 500 ng of RNA was used for cDNA synthesis (qScript cDNA Supermix, Quanta) and real-time quantitative PCR was performed (iTaq Universal SYBR Green Supermix, Bio-Rad) using CFX96 Touch Real-Time PCR Detection System (Bio-Rad). Each sample was run in triplicate wells, mean Ct values were obtained, and relative quantification of expression was calculated using the ΔΔCT method. Each gene was assayed with at least three biological replicates and all data was normalized to *Gapdh* in wild-type animals. Primer sequences are listed inin Supplementary Table 1.

### Western Blot

Eight-week old testes were de-tunicated and placed in T-PER lysis buffer (ThermoScientific, #78510) with protease inhibitor added (Sigma, #11836170001). Tissue was homogenized and spun down at 14,000 x g at 4°C for 10min to pellet cell debris. The supernatant was moved to a fresh tube and loading buffer (0.2M Tris HCl pH 6.8/6% SDS/30% glycerol/bromophenol blue/10% beta mercaptoethanol) was added. Samples were heat denatured at 95°C for 5min and cooled. Samples were loaded into a precast polyacrylamide gel (Bio-Rad, #4561096) Running buffer was 0.15M Tris/1M glycine/0.02M SDS. Electrotransfer was onto an activated nitrocellulose membrane. After protein transfer, the blot was washed in TBS-Tween (0.01%) and blocked with 5% milk/TBST for 1hr at RT. To probe for SPO11, the anti-SPO11 antibody (gift from S.K. or Millipore Sigma, #MABE1167) was diluted 1:500 in blocking buffer and the membrane was incubated at 4°C for overnight. Next day, the blot was washed 3×10min with TBST, then probed with 1:2000 HRP-conjugated antimouse secondary antibody at RT for 1hr. The blot was washed 3×10min with TBST, then HRP substrate was added (Millipore, #WBLUR0100) and incubated for 3min at RT, then scanned.

