## Supplemental Figs and Table for "A segregating human allele of *SPO11* modeled in mice disrupts timing and amounts of meiotic recombination, causing oligospermia and a decreased ovarian reserve"

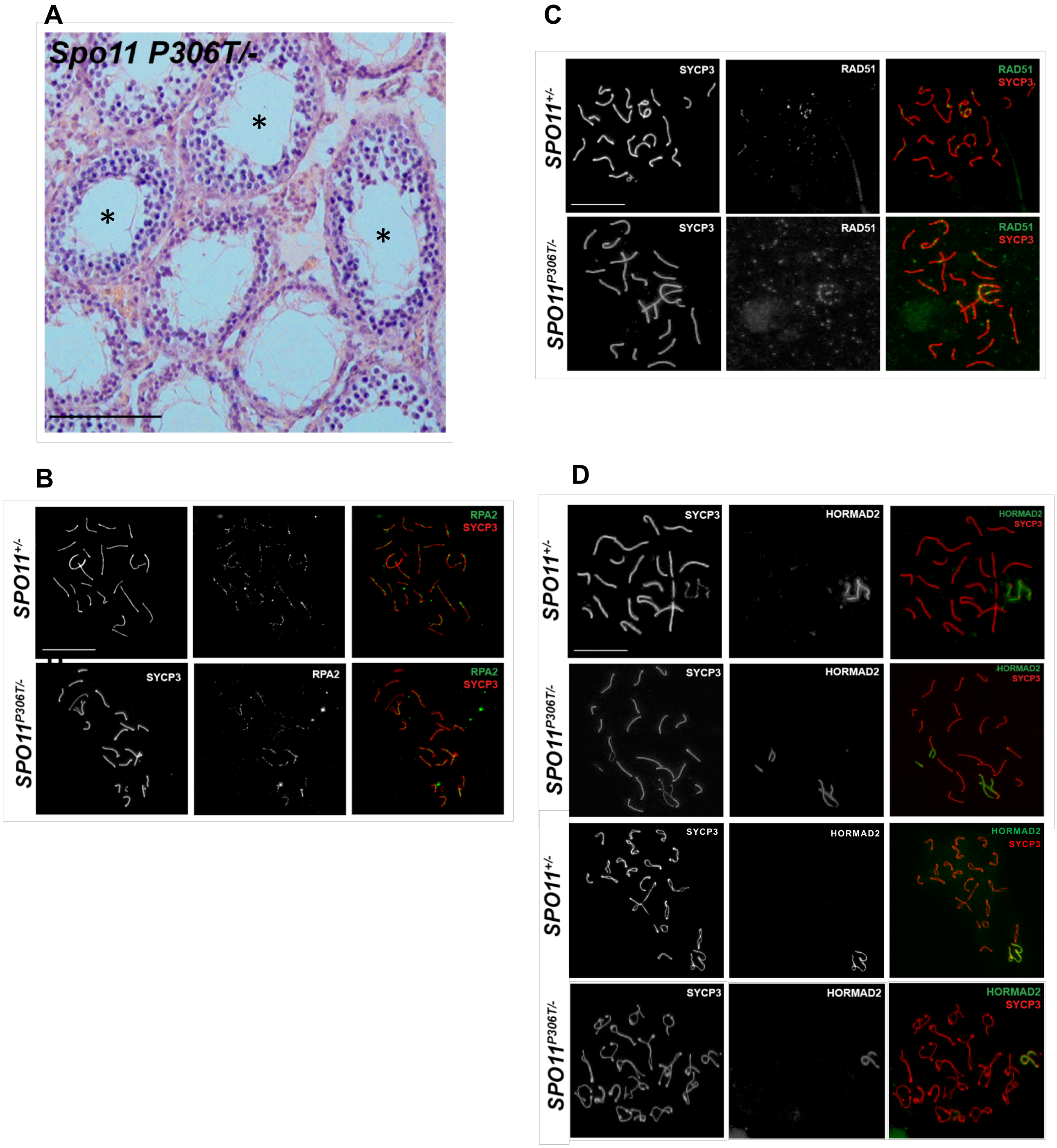


**Supplementary Figure 1.** *Spo11$^{P306T/-}$* males and females are severely germ cell depleted, and spermatocytes have drastically reduced meiotic DSBs. **(A)** H&E stained testis cross-section from an eight-week old *Spo11$^{P306T/-}$* male. Size bar = 75µm. Notice postmeiotic germ cells in all tubules. The tubules with asterisks appear to contain arrested spermatocytes. (**B, C)** Representative pachytene spermatocyte surface spread immunolabeled with RPA2 or RAD51 plus SYCP3. Quantification is plotted in Fig. 4. **(D)** Representative spermatocyte chromosome spreads immunolabeled with SYCP3 and HORMAD2 at pachynema (top two rows) and diplonema (bottom two rows). Whereas the only the XY body labels with HORMAD2, indicating asynapsed regions, there are also autosomes asynapsed at pachynema. Normal diplotene nuclei were also observed.

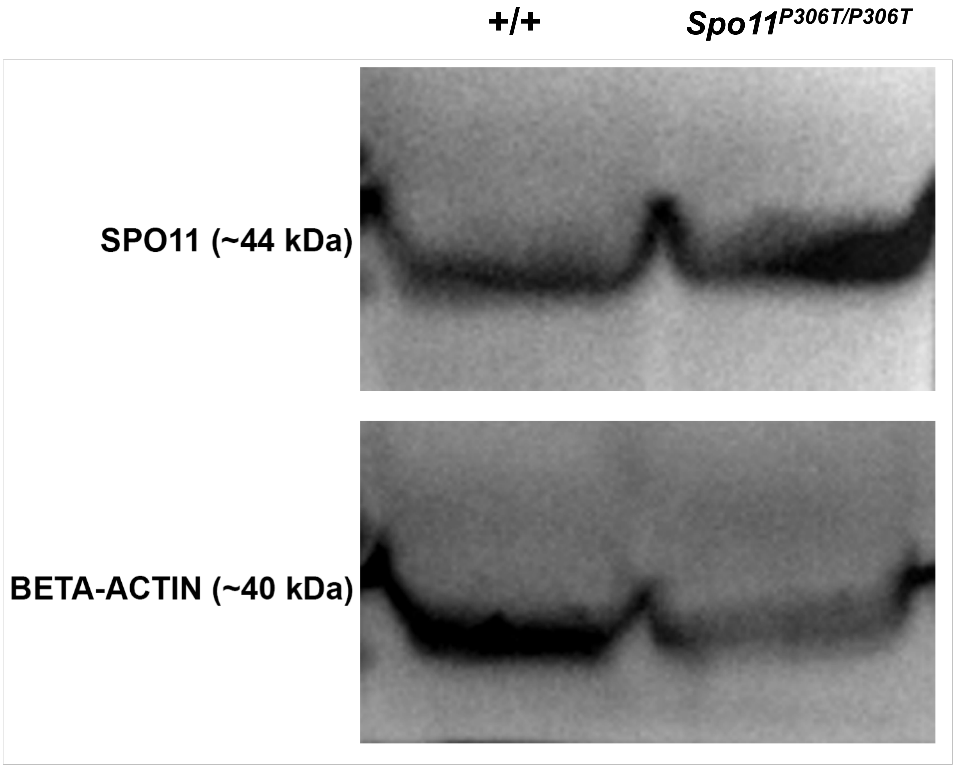


**Supplemental Figure 2.** Western blot of whole testis protein probed with anti-SPO11 antibody.

### Supplementary Table 1

| Purpose | Name | Sequence (5'-3') | Amplicon | RE Digest | Citation |
|---|---|---|---|---|---|
| Genotyping SPO11 P306T | Forward | GTTTCACCATGCCTGGACTT | 309 bp | HaeIII WT = 247bp, 62bp P306T = 125bp, 122bp, 62bp | |
| | Reverse | CCTGTAGCTCAGCCAGGTTC | | | |
| Genotyping SPO11 null | Mutant Forward | CTGCAGGTTTGATGATTCTGT | 200 bp | N/A | Baudat et al. 2000 |
| | Mutant Reverse | CATCAGAAGCTGACTCTAGAG | | | |
| | WT Forward | GCAATGCTCATTCTGTGTTG | 200 bp | Yes/No | |
| | WT Reverse | GGCACTTTCAGCATACAGGA | | | |
| Genotyping CHEK2 null | Mutant Forward | CCAAAGAAGTCTCCGTTGCT | 157 bp | Yes/No | Hirao et al. 2002 |
| | Mutant Reverse | CAAATTAAGGGCCAGCTCATTC | | | |
| | WT Forward | CCTTATGTGGTACGCCCACT | 150 bp | Yes/No | |
| | WT Reverse | CCACCTCATCCAACCAGACT | | | |
| RT-qPCR: DDX3Y | Forward | GGGCGCTATATACCTCCTCAC | [redacted] | [redacted] | Royo et al. 2010 |
| | Reverse | TCCAAAACTGCTGTAGGCATC | [redacted] | [redacted] | |
| RT-qPCR: EIF2S3Y | Forward | AACTATGCTGAATGGGGCAG | [redacted] | [redacted] | Royo et al. 2010 |
| | Reverse | TAATTTCAATGGCAGCCAGG | [redacted] | [redacted] | |
| RT-qPCR: SRY | Forward | GAGAGCATGGAGGGCCATG | [redacted] | [redacted] | Zwingman et al. 1993 |
| | Reverse | GAGTACAGGTGTGCAGCTC | [redacted] | [redacted] | |
| RT-qPCR: UBE2B | Forward | CGCCCCATCTGAAAACAACA | [redacted] | [redacted] | Royo et al. 2010 |
| | Reverse | TGGGACTCCATCGATTCTGC | [redacted] | [redacted] | |
| RT-qPCR: USP9Y | Forward | ATGGCAGGTTGCACATTCAC | [redacted] | [redacted] | Royo et al. 2010 |
| | Reverse | CAGTCCATCTTGATCATTTGG | [redacted] | [redacted] | |
| RT-qPCR: ZFY1 | Forward | GCCAGTGCTCTCTTAAACCAA | [redacted] | [redacted] | Royo et al. 2010 |
| | Reverse | TGAGTACACAAAGTCCCAGCA | [redacted] | [redacted] | |
| RT-qPCR: ZFY1/2 | Forward | TGGATGAAGCATCTCCAGAA | [redacted] | [redacted] | Royo et al. 2010 |
| | Reverse | CCACCAGCATCTTCATCTCC | [redacted] | [redacted] | |
| RT-qPCR: KDM6A | Forward | TACAGGCTCAGTTGTGTAACCT | [redacted] | [redacted] | Jiang et al. 2013 |
| | Reverse | CTGCGGGAATTGGTAGGCTC | [redacted] | [redacted] | |
| RT-qPCR: MID1 | Forward | AGAGAAACACAGAACTGGAGACT | [redacted] | [redacted] | Nakamura et al. 2017 |
| | Reverse | CAGTTTGGCTTCTTGACGGG | [redacted] | [redacted] | |
| RT-qPCR: | Forward | CTCATCCTCATGTCTTCTCCG | [redacted] | [redacted] | |

### Supplementary Table 1

| | | | | |
|---|---|---|---|---|
| XIST | Reverse | GATTCCAGATAGACAGGCTGG | [redacted] | Kobayashi et al. 2006 |
| RT-qPCR: GAPDH | Forward | CTTTGTCAAGCTCATTTCCTGG | | Kartha et al. 2016 |
| | Reverse | TCTTGCTCAGTGTCCTTGC | | |
